# Unpacking the Allee effect: determining individual-level mechanisms that drive global population dynamics

**DOI:** 10.1101/774000

**Authors:** Nabil T. Fadai, Stuart T. Johnston, Matthew J. Simpson

## Abstract

We present a solid theoretical foundation for interpreting the origin of Allee effects by providing the missing link in understanding how local individual-based mechanisms translate to global population dynamics. Allee effects were originally proposed to describe population dynamics that cannot be explained by exponential and logistic growth models. However, standard methods often calibrate Allee effect models to match observed global population dynamics without providing any mechanistic insight. By introducing a stochastic individual-based model, with proliferation, death, and motility rates that depend on local density, we present a modelling framework that translates particular global Allee effects to specific individual-based mechanisms. Using data from ecology and cell biology, we unpack individual-level mechanisms implicit in an Allee effect model and provide simulation tools for others to repeat this analysis.

## 1 Introduction

Understanding biological population dynamics provides insight into whether a population will survive or become extinct. Salient features of these population dynamics, such as the growth rate and the maximum population density, can be captured using suitable mathematical modelling frameworks^1–12^. A common approach to model the density of a population, *C*(*t*), is to describe the *per-capita growth rate* ^1^. *The percapita growth rate, f*(*C*), can be used to specify the temporal evolution of population density as

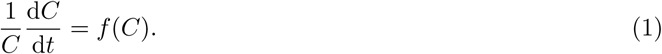

The most common mathematical descriptions of biological population dynamics are exponential and logistic growth models ^1^. Exponential growth is the simplest model, whereby *f*(*C*) is a positive constant (Fig. 1a). While the exponential growth model captures observed low-density population dynamics ^1–3;13^, exponential growth implies that the population will eventually become infinite. The logistic growth model (Fig. 1b) incorporates a linearly decreasing *f*(*C*) and is perhaps the most widely used model of biological and ecological population dynamics ^1–3;13^. This is because the logistic growth model captures two ubiquitous phenomena: (i) near-exponential growth at low density, and; (ii) a finite maximum density, termed the *carrying capacity* ^2–5;7;13;14^.

**Figure 1:**
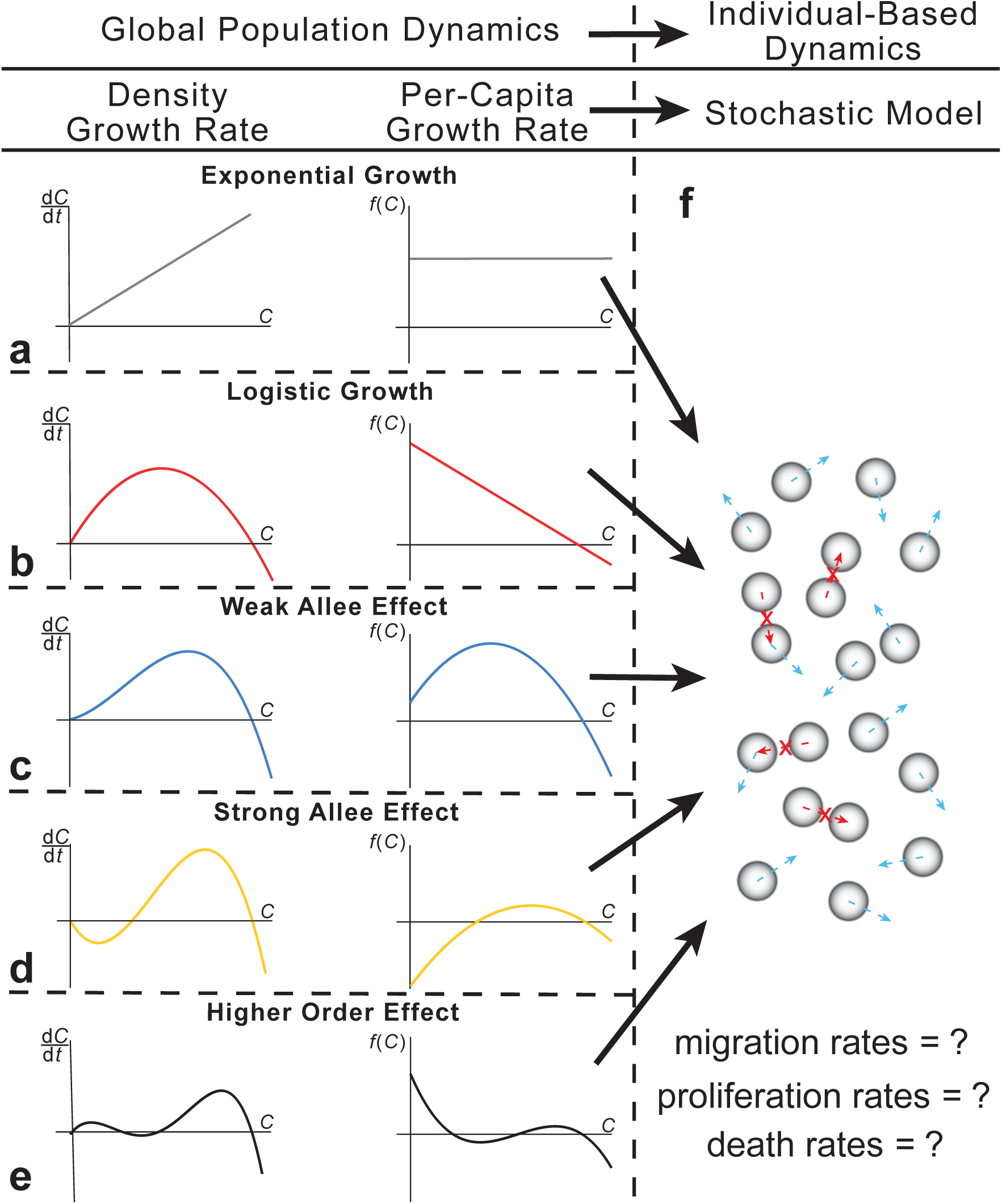
Relating global population-level models to an individual-based stochastic framework. A population can be described using one of two mathematical frameworks. In the first, global population models describe an averaged population density, *C*. These global population models are described in terms of the density growth rate, d*C/*d*t*, or the per-capita growth rate, *f*(*C*) = (1*/C*)(d*C/*d*t*). Common global models include (a) exponential, (b) logistic, (c) the Weak Allee effect, (d) the Strong Allee effect, and (e) higher-order modifications. In the second, a population can also be described using a stochastic individual-based model (f), where agents move, proliferate, and die based on stochastic rules (blue arrows). To ensure the stochastic model is as realistic as possible, crowding effects are incorporated by not permitting any event that would result in agent overlap (red crosses).

Classical exponential and logistic growth models rely on several key assumptions regarding the underlying biological mechanisms that drive population-level, or *global* dynamics ^6;8–11;15^. These assumptions include: all individuals survive at all densities ^1^, and the intrinstic growth and death rates are independent of density ^10^. However, populations have been observed to grow more slowly at low densities than predicted by the classical logistic model ^6^, while other populations undergo extinction below a threshold density ^7;15^. These observations are inconsistent with logistic and exponential models and modifications have been proposed to explain these observations ^2–7^.

The Allee effect (Fig. 1c,d) is a common modification of the logistic model that relaxes the assumption that all members of a population will survive. This model describes situations in biology where the per-capita growth rate is smaller, relative to logistic growth, at low population densities ^6;8–12;14;15^. The Allee effect is often discussed in the context of ecology and is relevant for describing the extinction of endangered species ^9;14^, population heterogeneity and structure ^16;17^, the impact of invasive species ^14;18^, and the reduction of biological fitness ^12;14^. While the Allee effect was first proposed in the ecology literature, more recent interest in the cell biology literature suggests that there is a growing awareness of the role of the Allee effect in populations of cells, including the eradication of cancer cells ^19;20^, growth rates of tumour cells ^2;3;20–22^, and cell migration and invasion assays ^10^. The Allee effect typically takes one of two forms, depending on the behaviour at low densities: (i) the *Strong Allee effect*, describing negative per-capita growth below some critical density threshold (Fig. 1d), resulting in the extinction of the population below this threshold, and; (ii) the *Weak Allee effect*, describing a reduced, but positive, per-capita growth rate at low densities (Fig. 1c).

Many studies incorporating Allee effects solely examine global information ^2;3;5;6;8;9;11;12;14;21^. Therefore, it is not obvious *a priori* how to determine an explicit form of an Allee effect from local, *individual-based* mechanisms. Alternatively, stochastic mathematical models can incorporate individual-level mechanisms to describe growing populations ^10;23;24^. These kind of individual-based model (IBM) simulation frameworks represent single members of the population as *agents* that, for example, move, proliferate, and die according to certain biologically-motivated stochastic rules.

IBMs are increasingly used to model population dynamics, partly because of technological advances making it possible to perform high-throughput cell biology experiments and collect large quantities of individual-level data. The task of choosing an appropriate model to capture and interpret experimentally-observed individual behaviour is a significant challenge ^25^. Typically, global population models are calibrated to experimental data to provide insight into the global features of the population, such as the carrying capacity or the low-density growth rate ^2;3;7^. However, this standard approach provides no insight into the underlying individual-level mechanisms. In contrast, employing an IBM allows us to describe both the local and global features of a population. While certain IBM frameworks proposed have been observed to produce Allee effects ^10;20;26;27^, the exact relationship between individual-based mechanisms and global per-capita growth rates is unclear. Consequently, the task of designing an IBM to describe *specific* global Allee effects, or higher-order effects (Fig. 1e), has never been considered.

In this work, we propose an IBM that incorporates motility, proliferation, and death processes in a population of individuals. This IBM incorporates crowding effects, whereby potential motility and proliferation events can only take place if agents do not overlap (Fig. 1f). By allowing individual-level mechanisms to depend on the density in a small neighbourhood surrounding an agent, the IBM is capable of describing a variety of per-capita growth rates. The continuum limit description of the IBM recovers the exact form of many Allee effect models for specific choices of IBM parameters. The main result is to pose and solve an inverse problem that can be described in the following way. Given a particular per-capita growth rate model, such as might be obtained by population-level experimental data, we determine which combinations of individual-level proliferation and death rates give rise to that particular scenario. This work provides the missing link in understanding between local, individual-based mechanisms and particular global outcomes, thereby providing a solid theoretical foundation for understanding and interpreting the mechanisms associated with Allee effects. We conclude by demonstrating how these new tools can be applied in practice by applying our modelling framework and solving the inverse problem to interpret data from both cell biology and ecology experiments. Additionally, interactive MATLAB applets, one of which determines IBM rates and simulations from a user-specified per-capita growth rate, and another that determines IBM rates from a user-specified choice of model fit to experimental data, are available to others to repeat this analysis (Code Availability).

## 2 Results

We consider a population of agents on a two-dimensional hexagonal lattice (Fig. 2). The IBM incorporates agent motility, proliferation, and death, where the individual-level rates vary with local density. All results in the main document consider the local density to be obtained by the six nearest neighbouring lattice sites (Fig. 2); additional results (Supplementary Information) show how our results generalise to larger neighbourhoods. Other types of regular lattices, including square lattices and three-dimensional cubic lattices, can also be used with the IBM framework.

**Figure 2:**
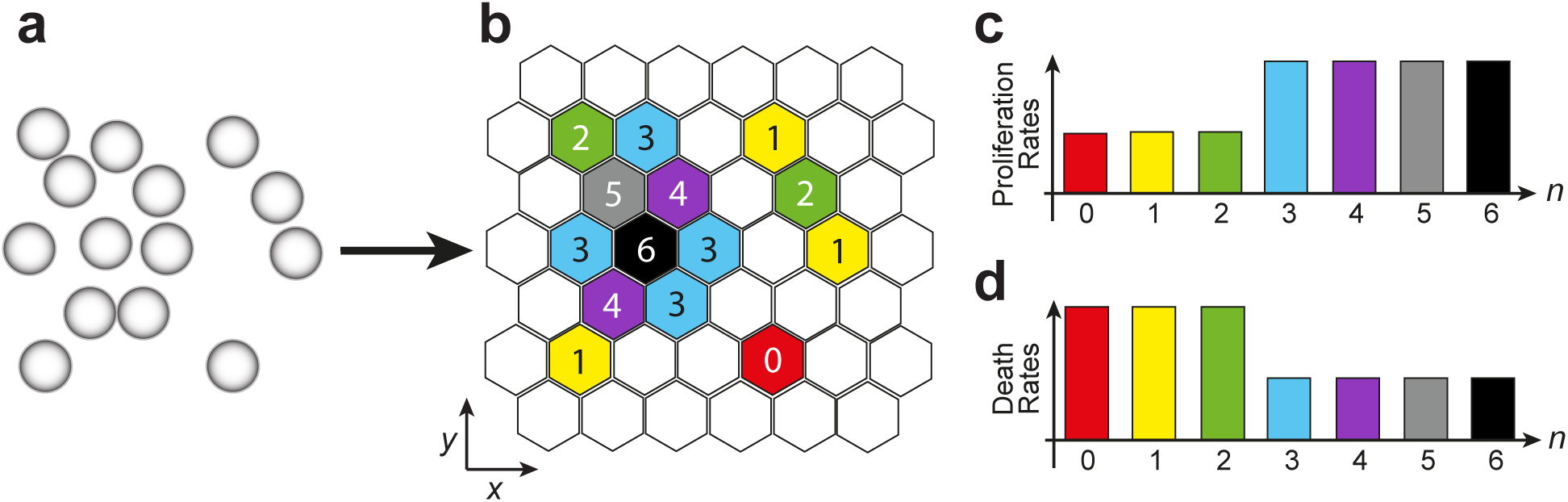
Agents on a hexagonal lattice with rates of proliferation and death that depend on the local density. Individuals within a population (a) are represented as agents on a two-dimensional hexagonal lattice (b). The corresponding proliferation and death rates of each agent are dependent on the local density of agents. The simplest measure of local density is the number of nearest neighbouring agents, *n*, as shown on the lattice. Here, *n* ranges from from *n* = 0 (red hexagon) to *n* = 6 (black hexagon). These differences in local density are used to specify how the attempted proliferation rates (c) and the death rates (d) depend on *n*. For example, in (b), there are three agents that each have one nearest neighbour (yellow hexagons). Each of these agents will have the same attempted proliferation and death rates, whose magnitudes are shown in (c,d) with yellow bars. We note that while an agent with six nearest neighbours (black hexagon) can attempt to proliferate (c, black bar), the net probability of this attempt being successful will always be zero. Nevertheless, an agent with six nearest neighbours can undergo a successful death event (d, black bar).

The main objective of this work is to determine how individual-level mechanisms are linked to various global Allee effects, which can be approached in two ways. The first approach is to demonstrate that this IBM framework gives rise to a variety of Allee effects, which we refer to as the *forward problem*, since the *input* of IBM parameters produces a certain global per-capita growth rate. The second approach is to determine which individual-level parameters give rise to a specific global per-capita rate. We refer to this as the *inverse problem*, as the inputs of the IBM parameters are unknown for a particular *output* per-capita rate. To highlight the insights obtained through this approach, we present strategies to link experimental data to various Allee effect models, which we interpret in terms of individual-level mechanisms.

### 2.1 The Individual-Based Model (IBM)

We perform non-dimensional simulations where the hexagonal lattice spacing is Δ. These simulations can be rescaled to match any particular application by rescaling Δ^23;28^. Each lattice site has position

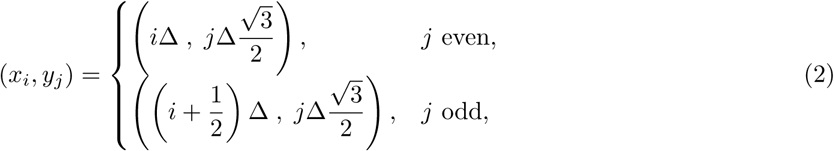

with *i* = 1, …, *I, j* = 1, …, *J*, and Δ = 1.

Crowding effects are important in both cell biology and ecology ^10;23;24;28–30^, so potential motility and proliferation events that would result in more than one agent per site are aborted. Agents attempt to undergo nearest neighbour motility events at rate *m*_*n*_ ≥ 0, proliferation at rate *p*_*n*_ ≥ 0, and death events at rate *d*_*n*_ ≥ 0 (Fig. 2c,d). Here, *n* ∈ {0, 1, …, 6} is the number of occupied nearest neighbour sites, providing a simple measure of the local density. While individuals *attempt* to undergo motility and proliferation events at a constant rate, the actual rate of *successful* events is density dependent, with the net rates being decreasing functions of density. This is the key feature of the model that gives rise to non-logistic phenomena.

For simplicity, we assume that all agents, regardless of their local density, have the same motility rate (*m*_*n*_ ≡ *m*). Furthermore, we choose *m* such that *p*_*n*_*/m* ≪ 1 and *d*_*n*_*/m* ≪ 1 for all *n*, since the characteristic timescale for motility is much shorter than the characteristic timescale for proliferation and death^31^. Agents are initially seeded on the lattice with a constant probability, representing spatially uniform initial conditions. Furthermore, we impose reflecting boundary conditions and, using a Gillespie approach ^32^, we simulate the number of agents as a function of time and space (Algorithm 1, Supplementary Information).

To compare data from the IBM with the global population description, we average data from the IBM using

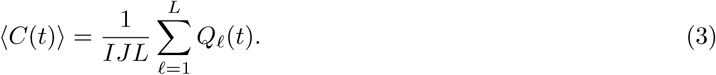

Here, *Q*_*ℓ*_(*t*) is the total number of agents on the lattice at time *t*, in the *ℓ*th identically-prepared realisation of the IBM. The total number of identically-prepared realisations is *L*; we choose *L* = 100 for the results presented in Figs. 3 and 4. A description of the numerical algorithm (Supplementary Information) and a MATLAB implementation of this algorithm are available (Code Availability).

**Figure 3:**
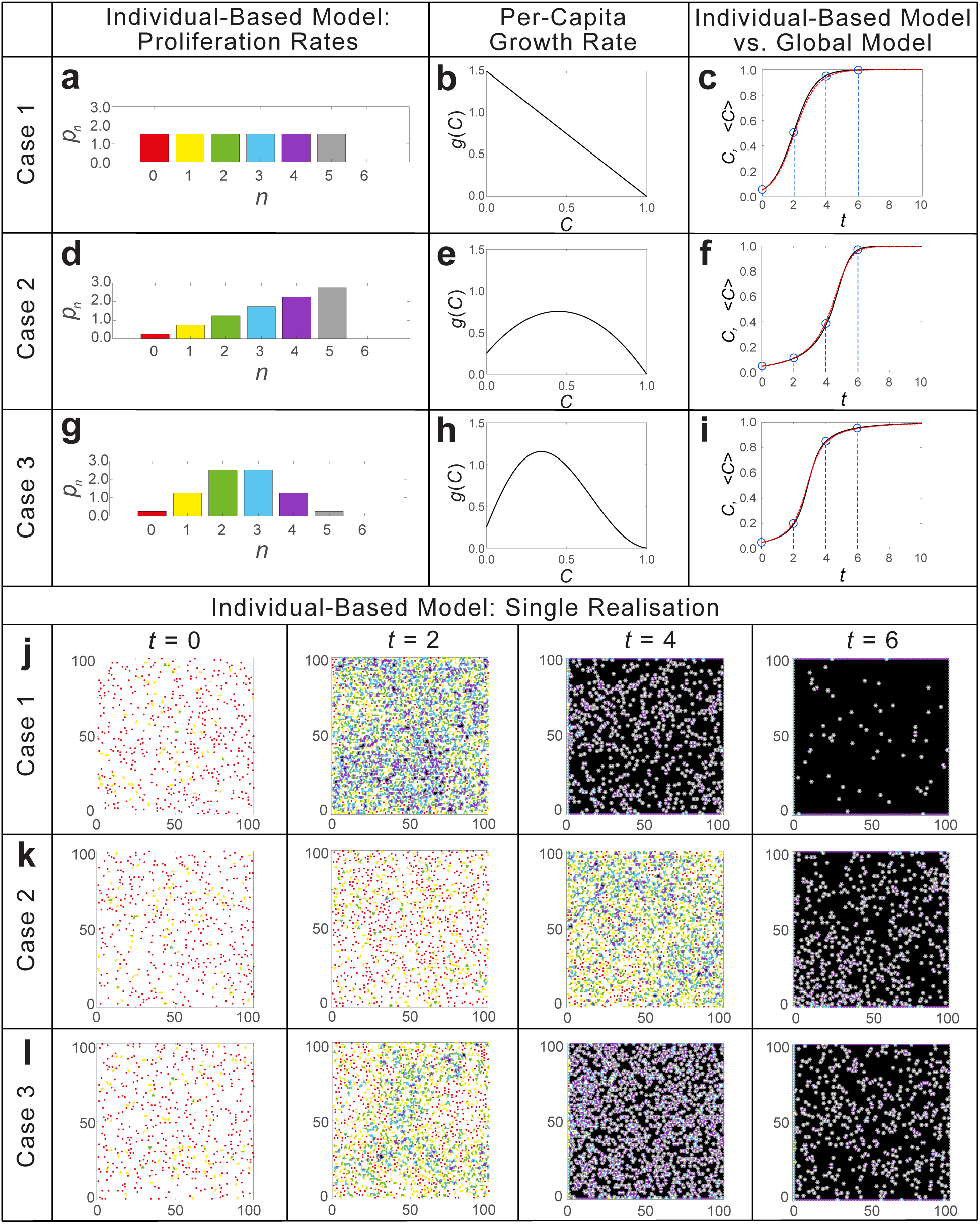
Comparison of data from the IBM with the solution of the corresponding global population description for different suites of proliferation rates. Simulations of the IBM are shown with three different familes of proliferation rates *p*_*n*_ (a,d,g), described in equation (10), along with the resulting per-capita rate of the global population model (b,e,h), described in equation (11). The three different families of proliferation rates are referred to as Case 1 (a–c,j), Case 2 (d–f,k), and Case 3 (g–i,l), respectively. (c,f,i) The solution of the global population description, *C*, is compared with averaged density data obtained by performing 100 identically-prepared realisations of the IBM to give ⟨*C*⟩, where the initial agent density is *C*(0) = 0.05. (j–l) Single realisations of the IBM, with the same colour scheme as in Fig. 2, are shown at *t* = 0, 2, 4, and 6, corresponding to the blue circles on the ⟨*C*⟩ curves. For each realisation of the IBM, we use a 100 × 115 hexagonal lattice, corresponding to the two-dimensional domain [1, 100] × [1, 100]. Here, *m* = 100 max(*p*_*n*_).

**Figure 4:**
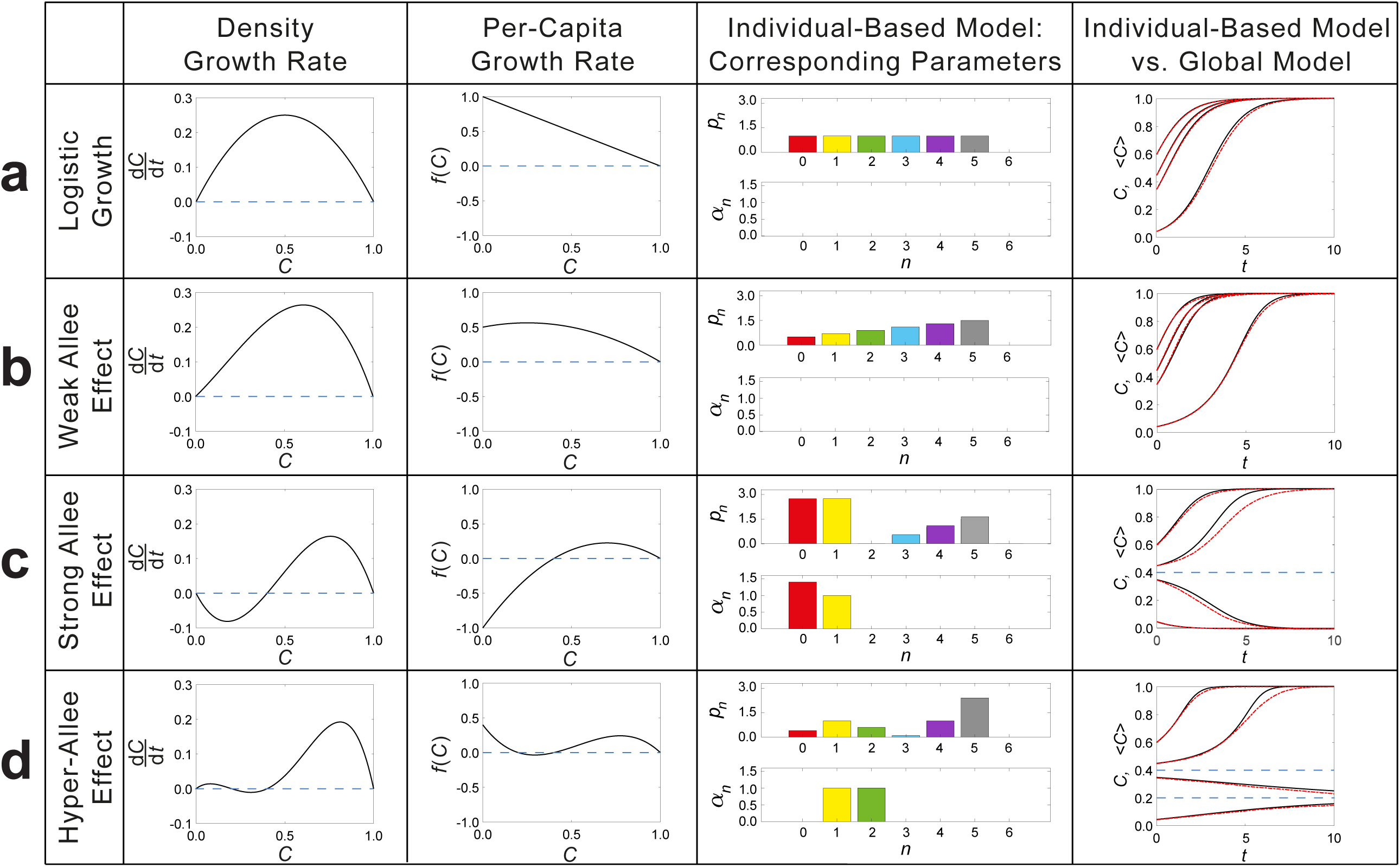
Summary of IBM mechanisms that correspond to particular choices of global per-capita growth rates. Four different choices of global density growth rates, along with their corresponding per-capita growth rates, *f*(*C*), are shown. (a) Logistic growth has *f*(*C*) = 1 − *C*; (b) the Weak Allee effect has *f*(*C*) = (*C* + 0.5)(1 − *C*); (c) the Strong Allee effect has *f*(*C*) = 2.5(*C* − 0.4)(1 − *C*), and; (d) the Hyper-Allee effect has *f*(*C*) = 5(*C* − 0.2)(*C* − 0.4)(1 − *C*). The corresponding IBM parameters are determined by solving equations (13) and (14) to determine the proliferation rates *p*_*n*_ and relative death rates *α*_*n*_. The solution of the global population description, *C*, is compared with averaged density data obtained by performing 100 identically-prepared realisations of the IBM to give ⟨*C*⟩, where four different initial agent densities are considered: *C*(0) = 0.05, 0.35, 0.45, and 0.6. For each realisiation of the IBM, we use a 100 × 115 hexagonal lattice while *m* = 100 max(*p*_*n*_).

### 2.2 Continuum limit

While the IBM allows us to visualise realistic-looking individual simulations of population dynamics, as well as to explicitly specify the individual-level behaviour of agents, it is convenient to derive a simpler mathematical description of the average behaviour of the IBM, called the *continuum limit description* ^10;28^. The continuum limit description gives us the ability to study global, deterministic, features of the IBM when the number of lattice sites is large, as well as to understand how individual-level differences translate into global outcomes.

Since the IBM employs spatially uniform initial conditions, the net flux of agents entering and leaving each lattice site due to motility events is, on average, zero. Therefore, spatial derivatives in the continuum limit will vanish, meaning that the continuum description of the average agent density, *C* ∈ [0, 1], is a function of time alone. Furthermore, we assume that the occupancy status of lattice sites is independent. This assumption, called the *mean-field approximation* ^10;23;28^, is mathematically convenient and is consistent with setting *m* ≫ *p*_*n*_ and *m* ≫ *d*_*n*_ ^23^. Our results confirm that the mean-field approximation is very accurate.

Using the mean-field approximation, we describe the local density of agents in terms of the average agent density. Specifically, for an agent to have a local density corresponding to *n* nearest neighbours, we require: (i) an agent to be present at the particular site; (ii) *n* nearest neighbour sites are occupied, and; (iii) the remaining (6 − *n*) nearest neighbour sites are vacant. These conditions give the density of agents with *n* nearest neighbours, *I*_*n*_ ∈ [0, 1], as

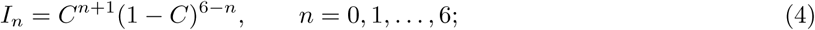

hence, *I*_*n*_ follows a binomial distribution. To determine how the global agent density evolves, we examine the time rate of change in expected agent density due to proliferation and death events, giving

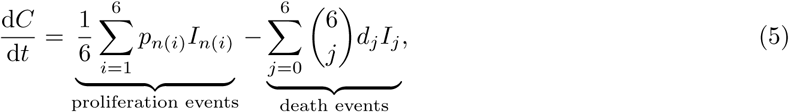

where *n*(*i*) is the number of nearest neighbours is each *neighbouring* agent *i*. The binomial coefficient, 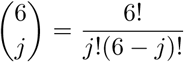, accounts for all possible configurations of an agent’s *j* nearest neighbours. Furthermore, by assuming that the agent configurations *I*_*n*_ follow a binomial distribution, we can write

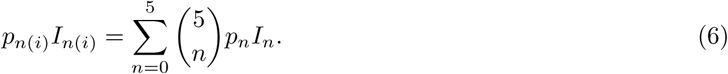

We omit the saturated configuration, *I*_6_ (black hexagon in Fig. 2), from this sum, since proliferation events require an empty neighbouring lattice site to produce a daughter agent.

By combining equations (5)–(6), we obtain the continuum limit description (i.e., the global population description) of *C*(*t*):

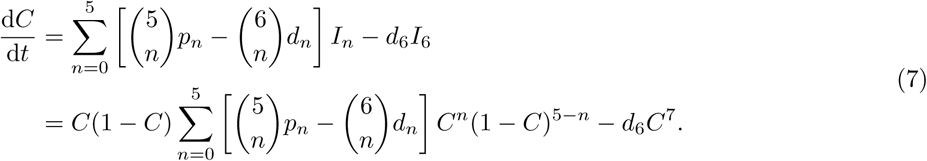

For convenience, we quantify the death rate as a fraction of the proliferation rate, i.e., *d*_*n*_ = *α*_*n*_*p*_*n*_, where *α*_*n*_ ≥ 0. With this substitution and some rearranging of terms in equation (7), we can write the evolution of the global agent density in terms of the per-capita growth rate, (1*/C*)(d*C/*d*t*):

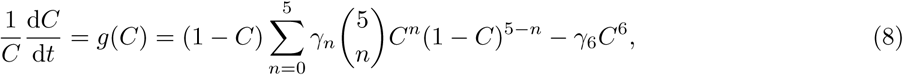

where

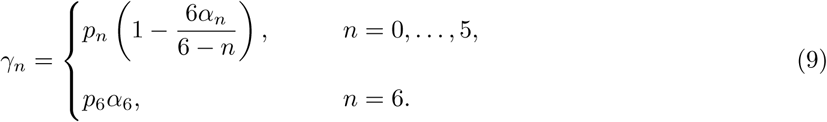

We deliberately refer to the per-capita growth rate associated with the continuum limit as *g*(*C*), and the per-capita growth rate prescribed via the global population behaviour as *f*(*C*). Later, we will show how to choose the IBM parameters such that *f*(*C*) ≡ *g*(*C*). Additionally, we note that equation (8) groups the 14 parameters (*p*_*n*_, *α*_*n*_) as 7 *linearly independent* parameters, *γ*_*n*_.

### 2.3 Allee effects arising from the IBM: the forward problem

We now demonstrate that the IBM framework gives rise to a rich variety of Allee effects. In the first instance, we set *α*_*n*_ = 0 for all *n*. Equation (9) implies that *γ*_*n*_ = *p*_*n*_ for 0 ≤ *n* ≤ 5 and *γ*_6_ = 0, such that agents only proliferate and move, and do not die. While this parameter regime is by no means a complete account of all possible parameter combinations, it serves to highlight that *g*(*C*), from equation (8), can give rise to a suite of Allee effects.

We consider three different choices of *p*_*n*_:

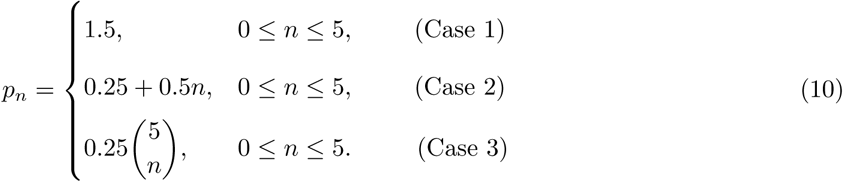

We note that in Cases 2 and 3, the range of proliferation rates varies by a factor of 10 as *n* varies from zero to five, demonstrating a significant range of proliferation rates within a single IBM realisation. Equation (8) gives,

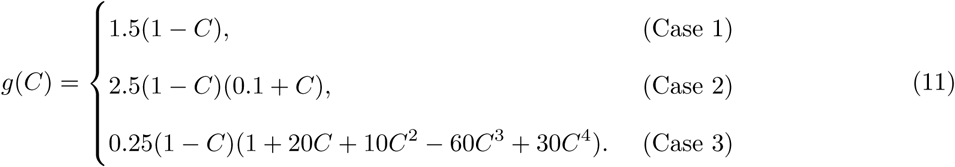

The global population density, *C*, can be determined by solving equation (8) with a specified initial condition, *C*(0).

While we can retrieve logistic growth from the continuum limit of the IBM (Case 1; Fig. 3), we can also obtain more complicated, nuanced, per-capita growth rates, including the Weak Allee effect (Case 2; Fig. 3) and completely novel Weak Allee-like per-capita growth rates never previously described (Case 3; Fig. 3). This analysis of three simple IBM parameter regimes suffices to show that this IBM framework is related to a large class of global population descriptions. Furthermore, in Fig. 3, as well as videos presented in the Supplementary Information, we show that averaged density data from the IBM, ⟨*C*⟩, agrees very well with *C*, confirming that the average agent density from the simulated IBM data is faithfully captured by the global population description. With these results, we now turn to the inverse problem to determine which individual-level rates describe global Allee effects.

### 2.4 Choosing IBM rates to match Allee effect models: the inverse problem

A standard practice for the application of an Allee-type model involves matching population-level experimental data to a particular continuum model, without any regard for the underlying individual-level mechanisms. Therefore, it is natural to ask, for a *given* per-capita growth rate, *f*(*C*), how do we choose the IBM parameters so that the per-capita growth rate determined from the continuum limit of the IBM, *g*(*C*) in equation (8), is identically *f*(*C*)?

Mathematically, we seek the parameters *γ*_*n*_ in equation (8) such that *g*(*C*) ≡ *f*(*C*). To do this, we first note that equation (8) is linear in *γ*_*n*_. Therefore, we can evaluate equation (8) at seven distinct points *C*_*i*_ ∈ {*C*_0_, *C*_1_, …, *C*_6_} and obtain a corresponding linear system in *γ*_*n*_. By denoting

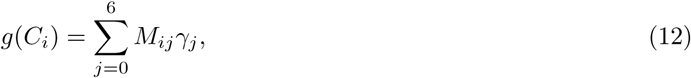

we obtain the linear system

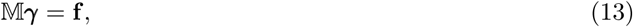

where the 7 × 7 matrix 𝕄 has entries [*M*_*ij*_], ***γ*** is the vector of parameter values [*γ*_*j*_], and **f** is the vector of function values [*f*(*C*_*i*_)]. We note that the polynomials appearing in (8), *C*^*n*^(1 − *C*)^6−*n*^, are linearly independent, so 𝕄 is never singular and the solution of this system is ***γ*** = 𝕄^−1^**f**. Furthermore, if *f*(*C*) is a polynomial of degree 6 or less and **f** ≠ **0**, this unique solution of ***γ*** provides the IBM parameters such that *g*(*C*) ≡ *f*(*C*). We discuss a method of determining the IBM parameters for other forms of *f*(*C*) in the Supplementary Information.

To prevent populations from becoming infinite, we require that *f*(*C*) will be non-positive when *C* = 1, so we impose *f*(1) ≤ 0. Noting that *g*(1) = −*γ*_6_ = −*p*_6_*α*_6_ ≤ 0, this implies that *γ*_6_ = −*f*(1) ≥ 0, which is consistent with the restriction that *p*_6_, *α*_6_ ≥ 0. Common per-capita rates and the corresponding IBM parameters are tabulated in the Supplementary Information.

Upon determining suitable parameters *γ*_*n*_ that match a specific *f*(*C*), we then consider choosing *p*_*n*_ and *α*_*n*_ from equation (9). However, since the birth and death rates (*p*_*n*_, *α*_*n*_) need to be determined solely from *γ*_*n*_, there are infinitely many different combinations of (*p*_*n*_, *α*_*n*_) that satisfy equation (9). If there is further information about either *p*_*n*_ or *α*_*n*_, such as additional experimental data, a unique estimate of (*p*_*n*_, *α*_*n*_) can be determined. However, in the absence of such data, we make a straightforward choice of parameter combinations:

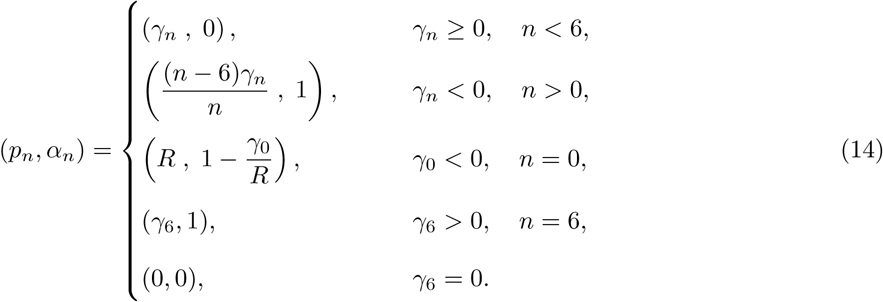

Here, 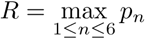 and only appears in the case when *f*(0) < 0. This choice of *R* provides a balance between minimising the relative death rate *α*_0_ > 1 while preventing *p*_0_ from dominating other proliferation rates. These parameters can be interpreted as proliferation-dominated behaviour when *γ*_*n*_ > 0 and death-dominated behaviour when *γ*_*n*_ < 0. Furthermore, having *γ*_6_ > 0 only occurs when *f*(1) < 0, implying that the death rate *d*_6_ = *p*_6_*α*_6_, shown in black in Fig. 2, must be strictly positive.

#### 2.4.1 Comparing the IBM with global population models

With this systematic method of determining the IBM proliferation and death rates that match various choices of *f*(*C*), we compare the average agent density determined by the IBM, ⟨*C*⟩, with its corresponding global population description, *C*. Results in Fig. 4 show that the IBM agent density agrees very well with the global population description, provided that the initial agent density, *C*(0), is sufficiently far away from any unstable equilibria of the global population description. An unstable equilibrium is a particular density *C*^∗^ such that d*C/*d*t* at this density, *C*^∗^*f*(*C*^∗^), is zero, but agent densities near *C*^∗^ diverge *away* from the equilibrium. For example, logistic growth (Fig. 4a) has an unstable equilibrium at *C*^∗^ = 0, whereas the Strong Allee effect (Fig. 4c) has an unstable equilibrium for a positive *C*^∗^ (Fig. 4c has *C*^∗^ = 0.4).

IBM simulations where the density is close to an unstable equilibrium are dominated by stochastic noise ^9;11;33^; therefore, we do not expect that ⟨*C*⟩ will agree with *C* in these cases. Indeed, this disagreement is clear in Fig. 4 with initial conditions in the Strong Allee effect and Hyper-Allee effect close to their unstable equilibria (see Fig. 4 caption for the explicit forms of these per-capita rates). Nevertheless, the IBM sufficiently captures the salient features of a large class of Allee-type dynamics with a suitable choice of proliferation rates, death rates, and initial conditions.

### 2.5 Mechanistic interpretation of experimental data

Per-capita growth rates describing population growth and extinction can match the global trends in experimental data ^2–7^, but fail to provide any insight at the individual-level scale. In contrast, the mathematical tools presented in this work provide the missing link that connects various global per-capita rates to specific individual-level mechanisms, via solving equation (13). Therefore, we can provide insight into individual-level behaviour from experimental data with the following approach. First, we choose a per-capita growth rate to match the global features of the experimental data. Second, the associated global model parameters are then fit to the data: for example, by minimising the least-square error between the population model and the experimental data. Last, we solve the inverse problem and determine the associated IBM parameters that give rise to the experimentally-observed global behaviour.

To highlight the insight possible from this approach, we consider two population-level data sets and provide previously hidden detail about individual-level behaviour ^6;7^. In Johnson et al. ^6^, BT-474 breast cancer cells are seeded at three initial densities in a 96-well plate, and cell proliferation is observed for 328 hours. Similar epithelial cell lines 20–30 µm in diameter have motility rates of about 4–9 h^−1^, while their proliferation rates are approximately 0.03–0.06 h^−1^ ^31^. Therefore, we assume that these BT-474 breast cancer cells have motility rates that are sufficiently large compared to both proliferation and death rates. As shown in the Supplementary Information, a motility rate that is ten times larger than the proliferation rate is sufficient for the well-mixed assumption of the IBM continuum limit to hold.

Model selection analysis in Johnson et al. ^6^ suggests that an Allee effect is required to describe this data. As there is no evidence of population extinction, we consider the Weak Allee effect with per-capita rate

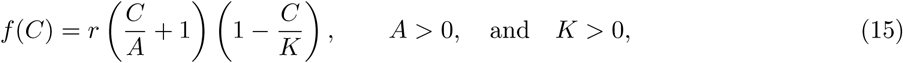

where *r* is the per-capita growth rate at *C* = 0 and *K* is the carrying capacity with units cells/image. We match this model to three experimental datasets simultaneously by minimising the total least-squares error (Fig. 5) and we determine the underlying individual-level mechanisms associated with equation (15) using equation (13). Specifically, the global Weak Allee parameters are determined to be *A* = 148 cells/image, *K* = 315 cells/image, and *r* = 0.00757 h^−1^. In Fig. 5b, we show that the IBM proliferation rates associated with this Weak Allee effect increase linearly with local density, as we observed when discussing the forward problem (Fig. 3). This is to be expected, since the Weak Allee effect also features a reduced growth rate at low densities. Furthermore, the individual-level rates increase by a factor of about three, which is significantly different to simpler models such as the logistic growth model, where the proliferation rates are independent of local density (see Fig. 4).

**Figure 5:**
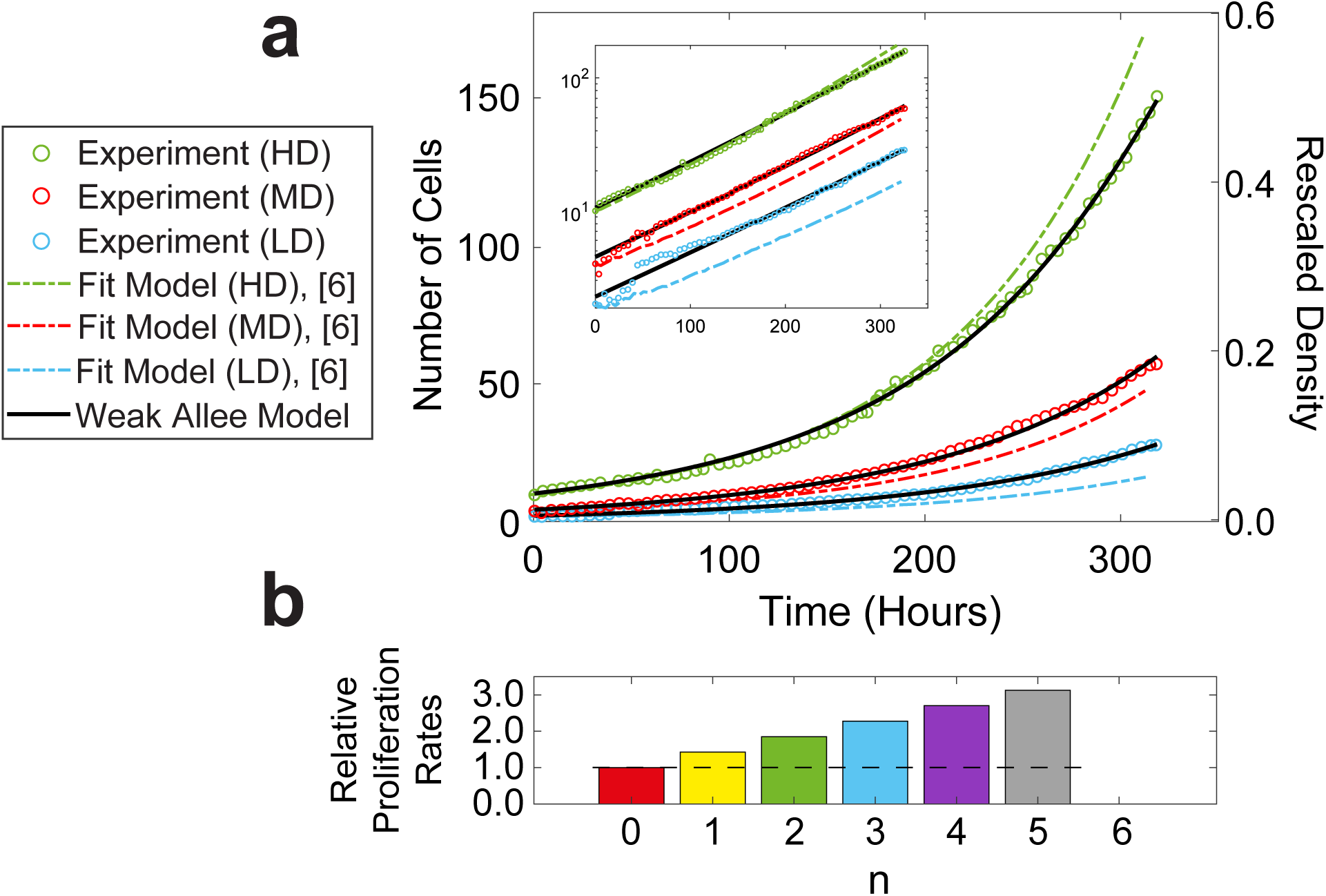
Number of BT-474 breast cancer cells compared to model predictions shown in Johnson et al. ^6^ and the Weak Allee effect. The number of BT-474 breast cancer cells are shown at low density initial conditions (LD), shown in blue circles, medium density initial conditions (MD), shown in red circles, and high density initial conditions (HD), shown in green circles, over the span of 328 hours ^6^. A semi-log plot of the data is shown in the inset of the figure to better distinguish the experimental data at low cell numbers. The three datasets can be fit to a modified Allee effect (dot-dash lines; parameters and model description shown in Johnson et al. ^6^). The Weak Allee effect (solid curves) is fit to minimise the combined least-square error of all experimental datasets shown in Johnson et al. ^6^. The Weak Allee parameters are determined to be *A* = 148, *K* = 315, and *r* = 0.00757, with fit initial conditions *c*_1_ = 2.27, *c*_2_ = 4.51, and *c*_3_ = 10.6. This parameter set yields the total least-squares error, combined over all three datasets, of 230, compared to the total least-squares error of 10900 using the model described in Johnson et al. ^6^. The rescaled density is the cell number data divided by *K*. Using this rescaled per-capita rate, we obtain the corresponding IBM parameters (b,c) by using equations (13) and (14). The proliferation rates (b) are shown relative to *p*_0_, shown in red, which has been rescaled by *r* so that *p*_0_ = 1. Unlike the logistic growth model (black dashed line), whose proliferation rates are independent of the local density *n*, the Weak Allee effect corresponds to proliferation rates that linearly increase with *n*. The magnitudes of the death rates are all zero.

As carrying capacity densities are often reported in units of cells/well, we note that *K* = 315 cells/image is equivalent to *K* = 33500 cells/well. To convert *K* from units of cells/image to units of cells/well, we make use of two key measurements: the size of the well, and the size of the image containing cells. The Trueline 96-well plate ^6^ has a well diameter of 6400 µm, while the largest region of an image shown in ^6^ that contains cells is approximately a 550 µm × 550 µm square. Therefore, we have that

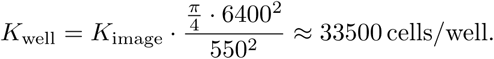

It is important to note that *K* = 33500 cells/well is simply the predicted carrying capacity density based on these low-cell number experiments at early time. Without any late time experimental data, where the density approaches the carrying capacity, it is impossible to predict the carrying capacity ^34^. Nevertheless, we infer the carrying capacity that minimises the least-squares error between model predictions and early-time experimental data. We conclude that the experimental observations presented in Johnson et al. ^6^ can be explained by allowing the attempted proliferation rates of BT-474 breast cancer cells to linearly increase as a function of local density. Without posing and solving the inverse problem described here, this individual-level insight is not possible.

To compare contiuum descriptions with experimental results that arise in ecology, we consider the datasets in Melica et al. ^7^, where the dynamics of the population density of *Aurelia aurita* polyps growing on oyster shells is measured. New *Aurelia aurita* polyps form via free-swimming propagules, which are released from the existing polyp into the surrounding water and land elsewhere on the oyster shell ^35^. Consequently, these polyps experience a faster motility rate, relative to proliferation and death rates, over the surface of the oyster shell. As such, the mean-field assumption inherent in our modelling framework is satisfied. Melica et al. ^7^ observe that a series of experiments where polyps are initially distributed at low density leads the population density evolve to some particular carrying capacity. Interestingly, when the same experiments are performed at a much higher initial density, the population decays to a different, higher, carrying capacity. This observation is explained in Melica et al. ^7^ by supposing the population dynamics are logistic, with the requirement that there are two different carrying capacities, despite the fact that the classic logistic model specifies a single carrying capacity only. Instead, we consider the “Hyper-Allee effect” model, Fig. 4(d), with per-capita rate

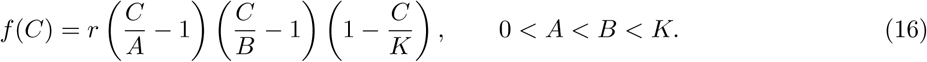

Equation (16) has the advantage that two stable equilibria, *A* and *K*, exist. While other cubic per-capita growth rates have been proposed ^36^, this work focuses on the underlying individual-level mechanisms giving rise to such a model. We fit this model to two experimental datasets simultaneously by minimising the combined least-squares error of both datasets. As seen in Fig. 6, the Hyper-Allee effect model agrees with both experimental datasets with only a change in the initial condition.

**Figure 6:**
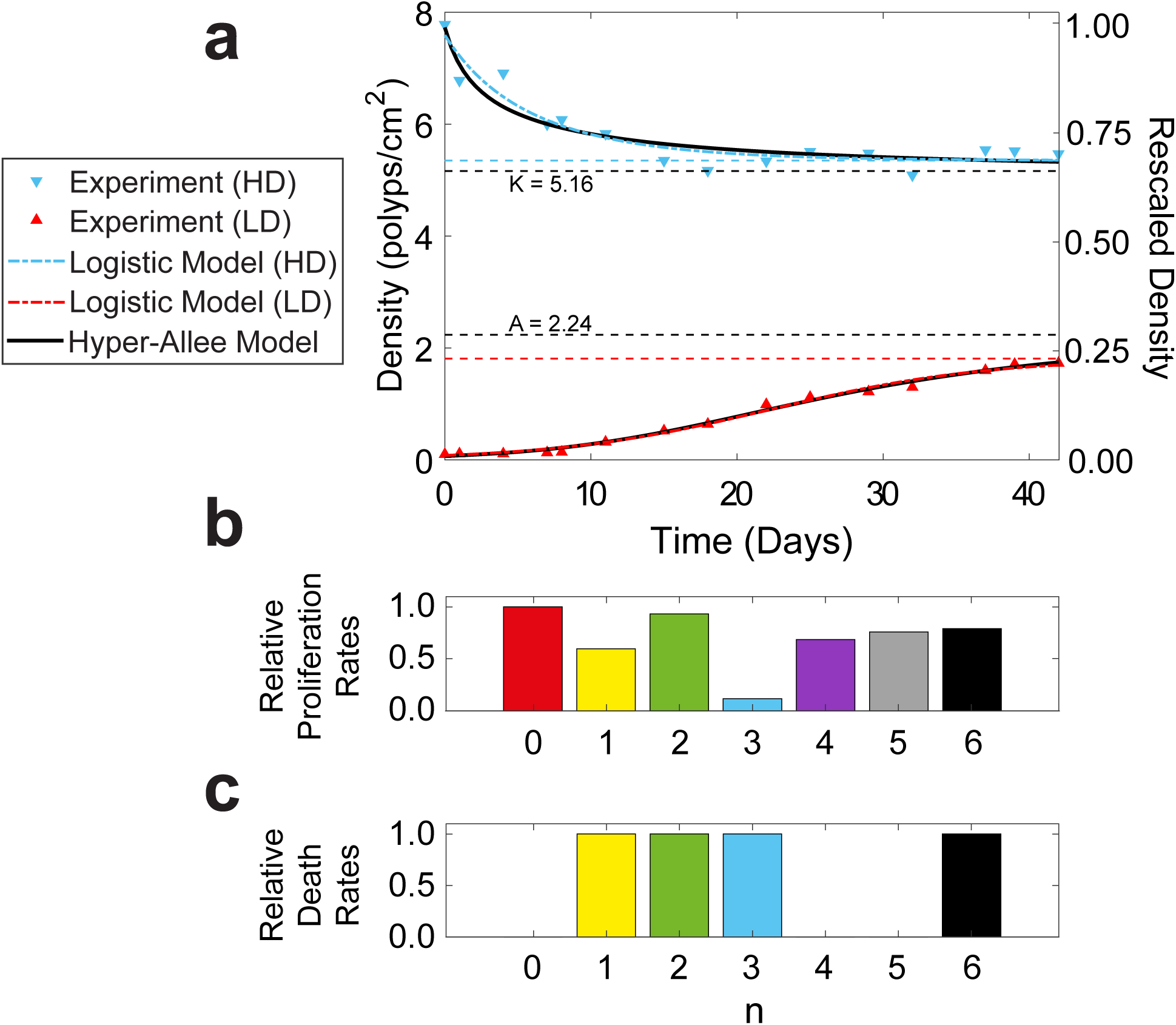
Density of *Aurelia aurita* polyps compared to model predictions of logistic growth and the Hyper-Allee effect. (a) The density of polyps are shown in low density treatments (LD), shown in red triangles, and high density treatments (HD), shown in blue triangles, over the span of 42 days. The two datasets can be fit to logistic growth separately (red and blue dot-dash lines; parameters are listed in Melica et al. ^7^), but cannot match both datasets simultaneously. The Hyper-Allee effect (equation (16), solid curves) is fit to minimise the combined least-square error of both expiremental datasets shown in Melica et al. ^7^. The Hyper-Allee parameters are determined to be *A* = 2.24, *B* = 4.69, *K* = 5.16, and *r* = 0.161, with two fit initial conditions *c*_1_ = 0.0691 and *c*_2_ = 7.73. The stable carrying capacities of both models are shown with dashed lines in the colours that correspond to their respective model. The rescaled density is the density data divided by the maximal experimentally observed polyp density, *c*_2_. Using this rescaled per-capita rate, we obtain the corresponding IBM parameters (b,c) by using equations (13) and (14). The attempted proliferation rates (b) are shown relative to *p*_0_, shown in red, which has been rescaled by *r* so that *p*_0_ = 1. The magnitudes of the death rates (c) are shown relative to their corresponding proliferation rates in (b). Since the rescaled density *C* = 1 is not an equilibrium point, the attempted proliferation and death rates when *n* = 6 (black bars) are non-zero.

To provide further insight about the population dynamics of this experiment, we determine the corresponding IBM parameters that give rise to this Hyper-Allee effect. We note that if *f*(*C*) is rescaled such that the largest recorded density point, the high-density initial condition, corresponds to *C* = 1 (Supplementary Information), then we must have *f*(1) < 0. Equation (13) implies that *γ*_6_ > 0, producing non-zero attempted proliferation and death rates when an individual has *n* = 6 nearest neighbours, shown in black bars in Fig. 6 b,c. However, the presence of death rates at various local densities (Fig. 6c) drives low density populations to the smaller carrying capacity *A*, while high density populations are driven to the larger carrying capacity *K*. We conclude that the experimental observations in Melica et al. ^7^, displaying the co-existence of two stable population densities, can be explained by varying the death rates with local density. In both examples, there is a clear need to use global population models more nuanced than logistic growth to capture the experimentally-observed features.

## 3 Discussion

Allee effects were first proposed to describe observed behaviour of populations that exhibit features unable to be explained by classical models, such as exponential and logistic growth. Such classical models rely on heavily simplifying assumptions, including all population densities being able to survive, and the growth and death rates being independent of the local density. The Allee effect relaxes these assumptions and modifies the logistic model by reducing the per-capita growth rate of a population to small or negative values at low densities, resulting in the potential extinction of a population below some critical density threshold. However, examination of Allee effects are nearly always performed at a global population scale alone. As such, unpacking the underlying, individual-level mechanisms that match a particular global Allee effect model has remained an open question in biology and ecology.

We demonstrate that by permitting proliferation and death rates in the IBM to vary with local density, we retrieve a large family of global population per-capita growth rates, including Allee effects. Furthermore, we propose a systematic method to determine individual-level mechanisms in the IBM that agree with a given particular per-capita growth rate model, such as might arise from experimental data. For example, once a per-capita growth rate is chosen to match the global features of experimental data, we can solve an associated linear system to determine the IBM parameters that give rise to the experimentally-observed global behaviour. While these IBM parameters cannot be uniquely identified without additional information, the underlying *net-growth* mechanisms in the IBM are uniquely identifiable. This method demonstrates that commonly used global per-capita growth rates, such as logistic growth, the Strong Allee effect, and the Weak Allee effect, can all be recovered with this IBM.

In this work, we also present strategies to connect experimental data to various global population models, which can in turn be linked to individual-level mechanisms. Specifically, we examine two experimental datasets arising in cell biology and ecology ^6;7^. In both cases, an Allee effect describes these datasets, and the global model parameters are not sensitive to initial conditions as suggested previously. Consequently, the IBM associated with these Allee effects not only provides better understanding of individual-level behaviour, but also provides insight into the limitations of commonly used per-capita rates. This IBM is useful for both theorists and experimentalists when analysing population dynamics; the modelling framework is simple to use and interpret, yet the insight and implications of the framework are broad and widely applicable.

By shifting to a modelling paradigm that involves the combination of an individual-based framework and its corresponding population dynamics, we provide insight into the behaviour of individuals from observed global behaviour. This insight provides a solid justification for the inclusion of more complex individual-level mechanisms to describe the salient features of population behaviour. The modelling approach proposed here provides a framework capable of unifying common global population descriptions with more complex population descriptions. Indeed, our results are not in conflict with the most common global population descriptions, including logistic growth and various forms of the Allee effect. Instead, our work highlights that by building a model from the individual-scale up, we systematically recover the basic underlying mechanisms that provide insight into whether a population will survive or become extinct.

### Code availability

All MATLAB codes are available at https://github.com/nfadai/Fadai_Allee2019.

### Data availability

All data are available upon request from the authors.

## Acknowledgements

This work is supported by the Australian Research Council (DP170100474, DE200100988). N.T.F. thanks Alex Browning (QUT) for help on creating the MATLAB applets. M.J.S. is also supported by the University of Canterbury Erskine Fellowship.

## Authors’ contributions

N.T.F. created the algorithm code and MATLAB applets, produced all figures, and carried out the analysis; M.J.S. and S.T.J. conceived, designed, supervised, and coordinated the study and helped draft the figures. N.T.F. wrote the paper, on which all other authors commented and revised. All authors gave final approval for publication.

## References

[1] Murray JD. Mathematical Biology I: An Introduction. Spring-Verlag; 2003.

[2] Sarapata EA, de Pillis LG. A comparison and catalog of intrinsic tumor growth Models. Bulletin of Mathematical Biology. 2014;76(8):2010–2024.

[3] Gerlee P. The model muddle: in search of tumor growth laws. Cancer Research. 2013;73(8):2407–2411.

[4] West GB, Brown JH, Enquist BJ. A general model for ontogenetic growth. Nature. 2001;413(6856):628.

[5] Tsoularis A, Wallace J. Analysis of logistic growth models. Mathematical Biosciences. 2002;179(1):21–55.

[6] Johnson KE, Howard G, Mo W, Strasser MK, Lima EABF, Huang S, et al. Cancer cell population growth kinetics at low densities deviate from the exponential growth model and suggest an Allee effect. PLoS Biology. 2019;17:e3000399.

[7] Melica V, Invernizzi S, Caristi G. Logistic density-dependent growth of an Aurelia aurita polyps population. Ecological Modelling. 2014;291:1–5.

[8] Stephens PA, Sutherland WJ, Freckleton RP. What is the Allee effect? Oikos. 1999;p. 185–190.

[9] Taylor CM, Hastings A. Allee effects in biological invasions. Ecology Letters. 2005;8(8):895–908.

[10] Johnston ST, Baker RE, McElwain DLS, Simpson MJ. Co-operation, competition and crowding: a discrete framework linking Allee kinetics, nonlinear diffusion, shocks and sharp-fronted travelling waves. Scientific Reports. 2017;7:42134.

[11] Dennis B. Allee effects in stochastic populations. Oikos. 2002;96(3):389–401.

[12] Lewis MA, Kareiva P. Allee dynamics and the spread of invading organisms. Theoretical Population Biology. 1993;43(2):141–158.

[13] Scott JG, Basanta D, Anderson ARA, Gerlee P. A mathematical model of tumour self-seeding reveals secondary metastatic deposits as drivers of primary tumour growth. Journal of The Royal Society Interface. 2013;10(82):20130011.

[14] Courchamp F, Clutton-Brock T, Grenfell B. Inverse density dependence and the Allee effect. Trends in Ecology & Evolution. 1999;14(10):405–410.

[15] Allee WC, Bowen ES. Studies in animal aggregations: mass protection against colloidal silver among goldfishes. Journal of Experimental Zoology. 1932;61(2):185–207.

[16] Ito H, Kakishima S, Uehara T, Morita S, Koyama T, Sota T, et al. Evolution of periodicity in periodical cicadas. Scientific Reports. 2015;5(14094).

[17] Larcombe MJ, Jordan GJ, Bryant D, Higgins SI. The dimensionality of niche space allows bounded and unbounded processes to jointly influence diversification. Nature Communications. 2018;9:4258.

[18] Seebens H, Blackburn TM, Dyer EE, Genovesi P, Hulme PE, Jeschke JM, et al. No saturation in the accumulation of alien species worldwide. Nature Communications. 2017;8:14435.

[19] Korolev KS, Xavier JB, Gore J. Turning ecology and evolution against cancer. Nature Reviews Cancer. 2014;14(5):371–380.

[20] Böttger K, Hatzikirou H, Voss-Böhme A, Cavalcanti-Adam EA, Herrero MA, Deutsch A. An emerging Allee effect is critical for tumor initiation and persistence. PLoS Computational Biology. 2015;11(9):e1004366.

[21] Axelrod R, Axelrod DE, Pienta KJ. Evolution of cooperation among tumor cells. Proceedings of the National Academy of Sciences. 2006;103(36):13474–13479.

[22] Lolas G, Bianchi A, Syrigos KN. Tumour-induced neoneurogenesis and perineural tumour growth: a mathematical approach. Scientific Reports. 2016;6:20684.

[23] Baker RE, Simpson MJ. Correcting mean-field approximations for birth-death-movement processes. Physical Review E. 2010;82(4).

[24] Fahse L, Wissel C, Grimm V. Reconciling classical and individual-based approaches in theoretical population ecology: a protocol for extracting population parameters from individual-based models. The American Naturalist. 1998;152(6):838–852.

[25] Browning AP, McCue SW, Simpson MJ. A Bayesian computational approach to explore the optimal duration of a cell proliferation assay. Bulletin of Mathematical Biology. 2017;79(8):1888–1906.

[26] Scott SM, Bodine EN, Yust A. An agent-based model of Santa Cruz island foxes (Urocyon littoralis santacruzae) which exhibits an Allee effect. Letters in Biomathematics. 2014;1(1):97–109.

[27] Colon C, Claessen D, Ghil M. Bifurcation analysis of an agent-based model for predator-prey interactions. Ecological Modelling. 2015;317:93–106.

[28] Jin W, Penington CJ, McCue SW, Simpson MJ. Stochastic simulation tools and continuum models for describing two-dimensional collective cell spreading with universal growth functions. Physical Biology. 2016;13(5):056003.

[29] Boukal DS, Berec L. Single-species models of the Allee effect: extinction boundaries, sex ratios and mate encounters. Journal of Theoretical Biology. 2002;218(3):375–394.

[30] Tobin PC, Bjørnstad ON. Spatial dynamics and cross-correlation in a transient predator–prey system. Journal of Animal Ecology. 2003;72(3):460–467.

[31] Simpson MJ, Landman KA, Hughes BD. Cell invasion with proliferation mechanisms motivated by time-lapse data. Physica A: Statistical Mechanics and its Applications. 2010;389(18):3779–90.

[32] Gillespie DT. Exact stochastic simulation of coupled chemical reactions. The Journal of Physical Chemistry. 1977;81(25):2340–2361.

[33] Drake JM. Allee effects and the risk of biological invasion. Risk Analysis: An International Journal. 2004;24(4):795–802.

[34] Warne DJ, Baker RE, Simpson MJ. Optimal quantification of contact inhibition in cell populations. Biophysical Journal. 2017;113(9):1920–4.

[35] Melica V. Influence of substrate availability on the population growth and on asexual reproduction strategies of *Aurelia* sp. polyps. Masters thesis; University of Trieste, 2013.

[36] Ludwig D, Jones DD, Holling CS. Qualitative analysis of insect outbreak systems: the spruce budworm and forest. The Journal of Animal Ecology. 1978;47(1):315–332.

