## Supplementary Information for "Unpacking the Allee effect: determining individual-level mechanisms that drive global population dynamics"

#### Contents

**S1 IBM Algorithm** **2**

**S2 Choosing agent motility-to-proliferation ratios** **3**

**S3 Explicit solutions of IBM rates to match common per-capita rates** **4**

**S4 Determining IBM rates for higher-order per-capita rates** **5**

**S5 Rescaling experimental data to agree with IBM rates** **6**

**S6 Extending the spatial template** **7**

### S1 IBM Algorithm

**Algorithm 1:** Pseudo-code for a single realisation of the IBM.

Create a two-dimensional $I \times J$ hexagonal lattice with some user-specified placement of agents; the total lattice sites is $IJ$;
Set $t = 0$;
**while** $t < t_{\text{end}}$ and $Q(t) < IJ$ **do**
    Randomly choose an agent and determine the number of nearest neighbours $n$;
    Calculate propensity function $a(t) := (m + p_n + d_n)Q(t)$;
    Calculate the following random variables, uniformly distributed on $[0, 1]$: $\beta_1, \beta_2$;
    Calculate time step $\tau = -\log_{\text{e}} \beta_1 / a(t)$;
    $t = t + \tau$;
    $Q(t) = Q(t - \tau)$;
    Calculate $R = a(t)\beta_2$;
    **if** $R \leq mQ(t)$ **then**
        Randomly choose adjacent hexagonal direction;
        **if** lattice site is unoccupied **then**
            Move agent to chosen site;
        **else**
            Nothing happens;
    **else if** $R \in (mQ(t), (m + p_n)Q(t)]$ **then**
        Randomly choose adjacent hexagonal direction;
        **if** lattice site is unoccupied **then**
            Add agent to chosen site;
            $Q(t) = Q(t) + 1$;
        **else**
            Nothing happens;
    **else**
        Remove agent;
        $Q(t) = Q(t) - 1$;

#### S2 Choosing agent motility-to-proliferation ratios

The continuum limit of the IBM assumes that individuals are well-mixed and fast moving over the lattice, since the proliferation and death rates are much smaller than migration. One natural question to ask is how large the motility rate has to be, relative to the proliferation rate, for the well-mixed assumption to how. To answer this question, we consider the motility-to-proliferation ratio

$$\mu = \frac{m}{\max(p_n)}, \tag{S1}$$

where $m$ is the motility rate independent of nearest neighbours and $p_n$ is the proliferation rate with $n$ nearest neighbours. As the well-mixed assumption will become increasingly valid as $\mu \to \infty$, we expect that the IBM increasingly matches the global population description for larger values of $\mu$. However, Fig. S1 shows that values of $\mu$ larger than 10 do not drastically alter the agreement between the IBM and the global population description. Consequently, we expect that populations of individuals which move at least 10 times more often than they proliferate can be described using the IBM.

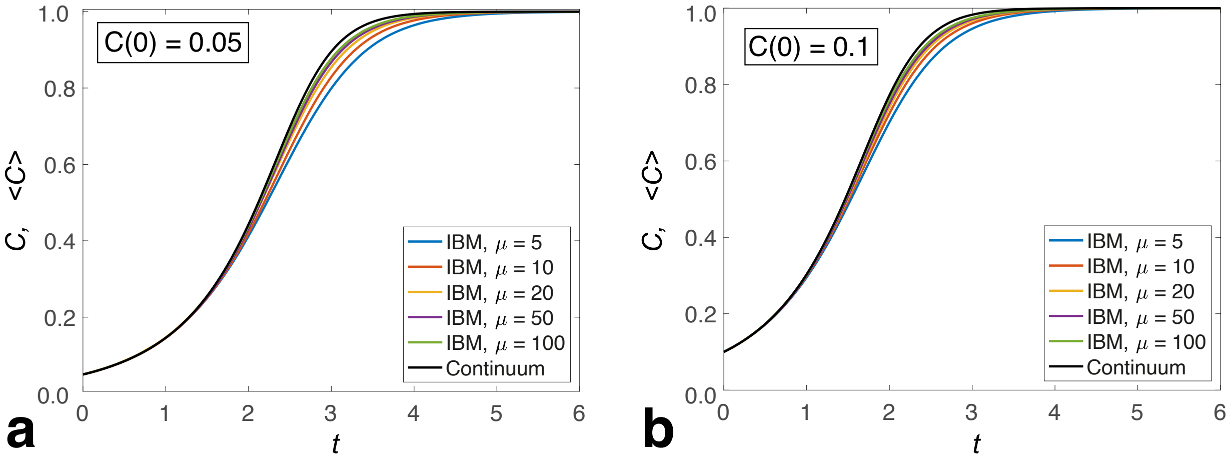


Figure S1: **Comparison of the Weak Allee effect with the average agent density of the IBM, for increasing motility-to-proliferation ratio $\mu$.** The Weak Allee global per-capita rate is $f(C) = 2C(1-C)(C+0.5)$ and the corresponding IBM parameters are $p_n = 1+0.4n, 0 \leq n \leq 5$ and $\alpha_n = 0$. The solution of the global population description is compared with averaged density data obtained by performing 100 identically-prepared realisations of the IBM to give $\langle C(t) \rangle$, where the initial agent density is (a) $C(0) = 0.05$ (b) $C(0) = 0.1$. For each realisiation of the IBM, we use a $100 \times 115$ hexagonal lattice while $m = \mu \max(p_n)$.

#### S3 Explicit solutions of IBM rates to match common per-capita rates

From the analysis in the "inverse problem" section, we now know that for any per-capita growth rate, $f(C)$, that is a polynomial of degree 6 or less with $f(1) \leq 0$, the solution $\gamma = \mathbb{M}^{-1}\mathbf{f}$ recovers $f(C)$ identically for a suitable choice of interpolation points $C_i$ via equation (2.12). With this, we examine four different per-capita growth rates:

- Logistic Growth, $f(C) = A - C, \qquad 0 < A \leq 1,$
- the Weak Allee effect, $f(C) = (A - C)(C + B), \qquad 0 < A \leq 1,\ B > 0,$
- the Strong Allee effect, $f(C) = (A - C)(C - B), \qquad 0 \leq B < A \leq 1,$
- a "Hyper-Allee effect" with two stable equilibria, $f(C) = (A - C)(C - B_1)(C - B_2)$, $0 \leq B_{1,2} < A \leq 1$.

All four per-capita growth rates have the property that $f(1) \leq 0$ for suitable choices of $A, B_1$, and $B_2$. Using equation (2.12), we can calculate the IBM parameter groupings $\gamma_n$ in equation (2.8), which are listed in Table S1.

| Effect Name | Logistic | Weak Allee | Strong Allee | Hyper-Allee |
|---|---|---|---|---|
| $\frac{\mathrm{d}C}{\mathrm{d}t}$ | $C(A-C)$ | $C(A-C)(C+B)$ | $C(A-C)(C-B)$ | $C(A-C)(C-B_1)(C-B_2)$ |
| $\frac{1}{C}\frac{\mathrm{d}C}{\mathrm{d}t}$ | $A-C$ | $(A-C)(C+B)$ | $(A-C)(C-B)$ | $(A-C)(C-B_1)(C-B_2)$ |
| $\gamma_0$ | $A$ | $AB$ | $-AB$ | $AB_1B_2$ |
| $5\gamma_1$ | $6A-1$ | $A-B+6AB$ | $A+B-6AB$ | $6AB_1B_2-(AB_1+AB_2+B_1B_2)$ |
| $10\gamma_2$ | $15A-5$ | $15AB+5(A-B)-1$ | $-15AB+5(A+B)-1$ | $15AB_1B_2 - 5(AB_1+AB_2+B_1B_2)+A+B_1+B_2$ |
| $10\gamma_3$ | $20A-10$ | $20AB+10(A-B)-4$ | $-20AB+10(A+B)-4$ | $20AB_1B_2-10(AB_1+AB_2+B_1B_2)+4(A+B_1+B_2)-1$ |
| $5\gamma_4$ | $15A-10$ | $15AB+10(A-B)-6$ | $-15AB+10(A+B)-6$ | $15AB_1B_2-10(AB_1+AB_2+B_1B_2)+6(A+B_1+B_2)-3$ |
| $\gamma_5$ | $6A-5$ | $6AB+5(A-B)-4$ | $-6AB+5(A+B)-4$ | $6AB_1B_2-5(AB_1+AB_2+B_1B_2)+4(A+B_1+B_2)-3$ |
| $\gamma_6$ | $1-A$ | $1-AB-A+B$ | $1+AB-A-B$ | $1-AB_1B_2+AB_1+AB_2+B_1B_2-(A+B_1+B_2)$ |

Table S1: **Individual-level rates that correspond to common global per-capita growth rates.**

### S4 Determining IBM rates for higher-order per-capita rates

In the "inverse problem" section, we note that a given per-capita rate, $f(C)$, can be identically represented by IBM parameters, provided that $f(C)$ is a polynomial of degree 6 of less using equation (2.12). While this class of per-capita rates contains a large variety of proposed Allee effects, there are many other per-capita rates proposed that are not members of this class [1–3]. One way of addressing these other forms of $f(C)$ is to *approximate* the per-capita rate as a 6th degree polynomial. Many approximations can suitably approximate $f(C)$; we propose two simple methods. One method is to choose the interpolating polynomial that approximates $f(C)$ at seven uniformly-spaced points, $C_i$, in the interval $[0, 1]$, i.e. $C_i \in \{0, 1, ...6\}/6$.

Naturally, this requires that $f(C)$ is continuous in $[0, 1]$, which is not necessarily true for certain per-capita growth rates. For example, the Von Bertalanffy per-capita rate [1] has the canonical form

$$f(C) = C^{-1/3} - 1. \tag{S2}$$

Since this per-capita growth rate is undefined at $C = 0$, the uniformly-spaced interpolation approach discussed previously will not succeed. A similar issue appears when using the canonical Gompertz per-capita rate $f(C) = -\log(C)$ [1–3]. Instead, one could approximate this form of $f(C)$ by determining the 6th degree polynomial that minimises the least-squares error between $f(C)$ and the polynomial, over a user-specified set of function values. However, this approach is prone to numerical difficulties and suffers from poor extrapolation properties. For example, if we sampled the Von Bertalanffy per-capita rate at 100 points in the interval $[0.1, 1]$, the resulting "best-fit" 6th degree polynomial will not faithfully represent the growth curve on the interval $(0, 0.1)$. Contrastingly, sampling the Von Bertalanffy per-capita rate at 100 points in the interval $[0.001, 1]$ results in the "best-fit" 6th degree polynomial unable to capture the salient features of the per-capita growth on the entire interval. In our related interactive Matlab applet, the uniform interpolating method is used to approximate non-polynomial $f(C)$, with an error message displayed if $f(C)$ is undefined at $C = 0$ or if $f(1) > 0$.

### S5 Rescaling experimental data to agree with IBM rates

We showed in the main article that we are able to choose appropriate global population models to fit experimental datasets. However, it still remains to relate this fit model to appropriate IBM parameters determined via the inverse problem in equation (2.12). Crucially, we require that the fit model should be rescaled so that the rescaled density, $c = C/\kappa$, lies entirely in the interval $[0, 1]$. A suitable rescaling is to set the rescaling parameter, $\kappa$, to the maximum density of a population. If the maximum density is unknown, another suitable rescaling of the density is

$$c = \frac{C}{\kappa}, \quad \text{where} \quad \kappa = \max\{K, \max_j \max_k [x_j(t_k)]\}. \tag{S3}$$

Here, $K$ is the carrying capacity determined from the model fit to the experimental datasets $x_j(t_k)$. For either choice of $\kappa$, this density rescaling allows the global population description to be linked to the IBM in terms of total lattice occupancy. For example, the Hyper-Allee effect fit to the density of *Aurlia aurita* polyps has the rescaled per-capita rate

$$f(c) = \frac{r\kappa^3}{ABK}\left(c - \frac{A}{\kappa}\right)\left(c - \frac{B}{\kappa}\right)\left(\frac{K}{\kappa} - c\right). \tag{S4}$$

Equation (S4) allows us to determine the corresponding IBM rates, using equation (2.12).

#### S6 Extending the spatial template

The IBM assumes that the proliferation and death rates only depend on its six nearest neighbouring lattice sites. One possible extension is to extend the spatial domain for which nearby agent neighbours influence proliferation and death events. For example, if we extend the local density neighourhood distance of a particular agent from $\Delta$ to $r\Delta$, this agent can perceive a maximum of $\mathcal{Z}(r)$ neighbours. For $r \leq 6, \mathcal{Z}(r) = 3r(r+1)$ [4]; further modifications need to be made for larger $r$. Consequently, this means that our inverse problem, shown in equation (2.12), now employs a $[\mathcal{Z}(r)+1] \times [\mathcal{Z}(r)+1]$ matrix, as we have $\mathcal{Z}(r)+1$ parameters in the $\gamma$ vector. Furthermore, this implies that we must also evaluate $f(C)$ at $\mathcal{Z}(r)+1$ distinct points to complete the inverse problem.

As this larger region of local density will better improve the underlying mean-field approximation employed in deriving the IBM continuum limit [5], we expect that the IBM increasingly matches the global population description for larger values of $r$. However, Fig. S2 shows that extending the spatial template to larger values of $r$ does not drastically improve the agreement between the IBM and the global population description.

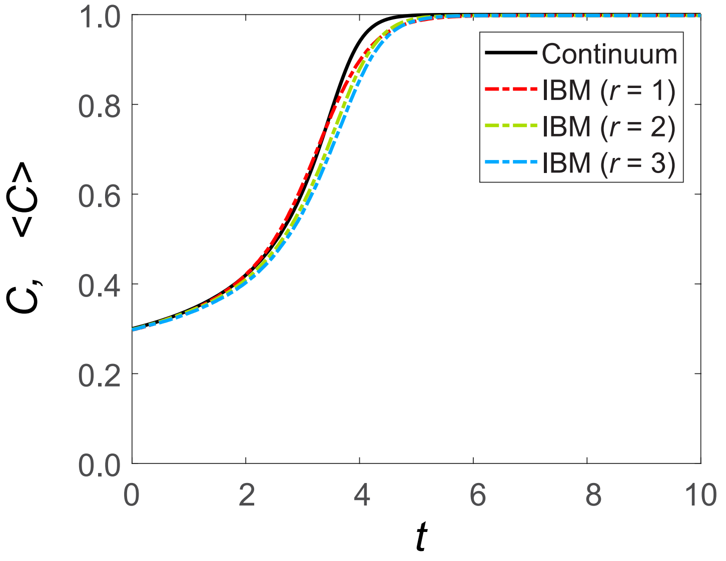


Figure S2: **Comparison of the Hyper-Allee effect with the average agent density of the IBM, for increasing radii of nearest neighbours $r$.** The Hyper-Allee global per-capita rate is $f(C) = 5C(1-C)(C-0.2)$ and the corresponding IBM parameters are determined by solving equations (2.12) and (2.13) to determine the proliferation rates $p_n$ and relative death rates $\alpha_n$. The solution of the global population description is compared with averaged density data obtained by performing 100 identically-prepared realisations of the IBM to give $\langle C(t) \rangle$, where the initial agent density is $C(0) = 0.3$. For each realisiation of the IBM, we use a $100 \times 115$ hexagonal lattice while $m = 100\max(p_n)$.

### References


[1] A. Tsoularis and J. Wallace, "Analysis of logistic growth models," *Mathematical Biosciences*, vol. 179, no. 1, pp. 21–55, 2002.

[2] P. Gerlee, "The model muddle: in search of tumor growth laws," *Cancer Research*, vol. 73, no. 8, pp. 2407–2411, 2013.

[3] E. A. Sarapata and L. G. de Pillis, "A comparison and catalog of intrinsic tumor growth models," *Bulletin of Mathematical Biology*, vol. 76, no. 8, pp. 2010–2024, 2014.

[4] W. Jin, C. J. Penington, S. W. McCue, and M. J. Simpson, "Stochastic simulation tools and continuum models for describing two-dimensional collective cell spreading with universal growth functions," *Physical Biology*, vol. 13, no. 5, p. 056003, 2016.

[5] N. T. Fadai, R. E. Baker, and M. J. Simpson, "Accurate and efficient discretizations for stochastic models providing near agent-based spatial resolution at low computational cost," *Journal of the Royal Society Interface*, vol. 16, no. 159, p. 20190421, 2019.
